# HONEY PRODUCTION IN FOREST AREAS: CHARACTERIZATION OF 5 HONEY SAMPLES FROM SOUTHERN CÔTE D’IVOIRE

**DOI:** 10.64898/2026.02.14.705139

**Authors:** Kouamé Koffi Félix, Assi Kaudjhis Chimène

## Abstract

The objective of this study is to determine the quality and define the different classes of honeys produced in the Ivorian forest region according to their pollen content. This involves the analysis of five honey samples from the sub-prefecture of Cechi. Four of the honey samples were wild-harvested, and one was from experimental beekeeping in the Cechi reserve. A total of 54 pollen taxa were identified. The most represented botanical families are: Fabaceae (9 species, or 16.67%), Apocynaceae, and Combretaceae, each with 5 species, or 9.27%. The pollen taxon richness of the honeys varies from 18 to 34 taxa. Most are polyfloral honeys, with the exception of the honey from the reserve, which contains 66.13% *Bridelia micrantha* pollen (Euphorbiaceae), a monofloral honey. These samples contain highly variable pollen content and fall into three categories of honey: honeys rich in pollen, honeys very rich in pollen, and honeys extremely rich in pollen, attesting to their high quality and natural origin.

## INTRODUCTION

Honey, produced from non-timber forests, is a naturally available source of sugar on the planet [1]. Honey is the best-known beehive product. It is made by honeybees from flower nectar or secretions from the living parts of plants or found on them, which the bees collect, transform, combine with their own specific substances, store, and allow to ripen in the honeycomb [2]. For its nutritional and therapeutic properties, honey has long been harvested by humans, first through gathering and then through beekeeping [3, 4, 5]. Honey owes all its properties to plants. According to N’guessan et al. [6], the therapeutic effects of honey are due to the flavonoids contained in plant compounds. Furthermore, honey owes its color to pigments such as carotenoids and flavonoids, which vary according to the geographical and floral origin [7].

Bees, while foraging on the flowers of the plants they visit, carry pollen, either intentionally or unintentionally, depending on the nutrients collected. This pollen is transported to the hive and ends up in the honey. In other words, inside the hive, pollen from the flowers foraged by the bees is mixed with the honey. Thus, melissopalynological analysis makes it possible to identify, based on the pollen content of the honey, the plants visited by the bees and the geographical origins of the honey. According to Tossou et al. [8] and Kouamé [4], melissopalynology is a complementary study to field inventories of melliferous plants; it provides a detailed and comprehensive understanding of the melliferous flora of an area. The study of melliferous plants is of great interest because it forms the basis for the objective evaluation of the quantitative and qualitative productivity of bees in different regions [9].

In Côte d’Ivoire, data on honey plants are limited and mostly generated in the central and northern parts of the country. This is evidenced by the work of Iritié et al. [10] on the identification of honey plants in the agroforestry zone of the Yamoussoukro Higher School of Agronomy; of Coulibaly et al. [11] on the potential of honey plants of interest for modern beekeeping in the Guinean savanna; of Kouassi et al. [12] on the diversity of honey plants in the Sub-Sudanese savanna; of Assi et al. [13] on the diversity of honey plants in the forest-savanna transition zone of Côte d’Ivoire in the department of Toumodi; and of Gnangouli bi et al. [14] on the pollen analysis of honeys from the Poro, Tcholo, Hambol, Bélier, and N’zi regions. A floristic inventory was carried out within the Yapi Daniel forest and extension in the southern forest region of Côte d’Ivoire [15]. The objective of this study is to determine the quality and different classes of honeys produced in the Ivorian forest region based on their pollen content.

## MATERIALS AND METHODS

### Study area

The study was conducted in Cechi, a sub-prefecture of the Agboville department located in the Agnéby-Tissa region of southeastern Côte d’Ivoire. This region belongs to the Guinean zone of Côte d’Ivoire [16], with a Guinean-type climate. The ecological zone is characterized by an average annual rainfall of 1585.35 mm and an average annual temperature of 26.72 °C [15]. The vegetation is a semi-deciduous humid dense forest, typical of the mesophilic zone of Côte d’Ivoire and the Guineo-Congolian center of floristic endemism. The population of the Cechi sub-prefecture is estimated at 22,779 inhabitants according to the 2021 general census of populations and habitats [17]. The main ethnic groups represented in the area are the Abbey, followed by the Krobou and migrant peoples from other parts of the country (Agny, Baoulé, Malinké) as well as from neighboring countries (Burkinabé, Mali, Guinea). These populations are mostly farmers, sometimes organized into associations or agricultural cooperatives.

### Sampling

Five (5) honey samples from localities within the Cechi sub-prefecture were used for this study. Four of the honey samples were collected from wild harvesting or traditional beekeeping practices, gathered from beekeepers in the localities of Allany (E1); Banguié 2 (E2); Mitichi (E4); N’guessanBlekro (E5), [18] and one honey sample from modern hives located in the reserve (E3) at BouaM’po.

### Honey Sample Processing

The technique used for processing honey samples is acetolysis. This method is based on that of the International Commission on Apicultural Botany described by Louveaux et al. [19], and has been adapted by several authors in their work to facilitate pollen observations [20]. For each well-homogenized honey sample, 10 g are weighed and dissolved in a beaker containing 20 ml of water acidified with sulfuric acid (5% H2SO4). The resulting solution is stirred with a magnetic stirrer until the honey is completely dissolved and centrifuged for 15 minutes at 3000 rpm. The supernatant is removed, and the first pellet obtained is rinsed with 10 ml of distilled water at 3000 rpm for 5 minutes. After rinsing, the supernatant is again removed to a depth of 1 to 2 cm from the pellet using a Pasteur pipette. Twenty microliters of the last pellet collected are transferred onto a microscope slide using a Pasteur pipette and placed in the oven at 40 °C for drying.

Once the preparation is dry, a drop of gelatinized glycerin is placed on top and carefully covered with a coverslip, ensuring no air bubbles penetrate. After removing any excess glycerin, the edges of the coverslip are sealed with clear nail polish to keep the preparation moist and free of air bubbles for microscopic examination.

### Observation and Identification of Pollen Grains

Pollen observations were performed using a binocular light microscope at 400x and 1000x magnification. Three slides were prepared for each honey sample. Pollen grains were identified using the Pollen Atlas of Côte d’Ivoire [21], the Pollen Atlas of West African Tropical Plants [22], the Pollen Atlas of Congo [23], the African Pollen Database [24], the online pollen database (https://data.oreme.org/palyno/palyno_gallery), and laboratory pollen data. Pollen characteristics considered included shape, size, exine ornamentation, and the presence or absence of apertures on the pollen grain. Pollen taxa were identified at the botanical family, genus, and species levels. Some pollen taxa are designated by the name of their genus or family followed by “type” (Ex: *Lannea-type*; *Cyperaceae-type*) according to the nomenclature proposed by the African Pollen Database by Vincens et al. [24] and adopted by some authors [25, 13]. Pollen taxa whose degradation of morphological structure does not facilitate identification are considered undifferentiated taxa and those that could not be identified as indeterminate or unidentified taxa (Ind).

### Qualitative Pollen Analysis

Qualitative pollen analysis consists of identifying and counting the pollen taxa present in a microscopic preparation [7]. It is carried out in two stages. The first involves the systematic identification of the pollen taxa present in the preparation; the second involves counting the pollen taxa.

Qualitative pollen analysis makes it possible to obtain the relative frequencies (RF) of the different pollen taxa and to obtain the pollen spectrum of the honey samples.

A pollen spectrum reflects the proportions of the various types of pollen identified in the honey. According to Comlan et al. [26], the sum of the pollen counts must reach at least 300 grains, and the total number of pollen taxa must be equal to or greater than 20 per sample. The pollen spectra of honeys are established by the relative frequencies of the pollen of the taxa encountered relative to the total number of pollen grains counted [27]. The relative frequency of pollen taxa is given by

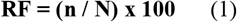

RF: relative frequency; n: number of pollen grains of a pollen taxon; N: total number of pollen grains counted in a preparation.

The assessment is based on Louveaux et al. [27] and Feller-Desmaly and Parent [28], who define four types of pollen grains according to their relative frequencies:

- Dominant pollen grains (RF ≥ 45%);
- Accompanying pollen grains (16 ≤ RF < 45%);
- Significant pollen grains (3 ≤ RF < 16%);
- Rare isolated pollen grains (RF < 3%).

The frequency of occurrence of pollen taxa is obtained by dividing the number of honey samples containing the taxon by the total number of honey samples analyzed.

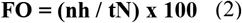

FO: frequency of occurrence of pollen taxa; nh: number of honey samples containing the taxon; tN: total number of honey samples analyzed.

This method establishes a correlation between the taxa inventoried in the honeys.

This association allows for the identification of the geographical origin of honeys, as it generally reflects the floristic composition of the vegetation in the environment [7]. The assessment is based on Louveaux et al. [19], who define four classes:

- Class 1: very frequent taxon (FO > 50%);
- Class 2: frequent taxon (20% < FO < 50%);
- Class 3: infrequent taxon (10% < FO < 20%);
- Class 4: rare taxon (FO < 10%).

### Quantitative pollen analysis

Quantitative pollen analysis allows for the evaluation of the amount of pollen contained in a unit weight of honey [30, 7]. It provides data on the pollen richness of honeys and the extraction method. The pollen content of honeys is expressed as the number of pollen grains per 10 g of honey, based on 20 µl of pollen preparation mounted between a slide and coverslip according to Louveaux et al. [19]. It is calculated using the following formula according to Erdtman [31]:

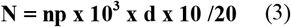

N: pollen content; np: number of pollen counted on a slide (20 µL); d: honey density.

The presentation of the results is based on that of Maurizio [32, 30], who classifies honeys into five categories according to their absolute pollen content (N). These are:

- Class I: N < 20,000 (honey with low pollen content);
- Class II: 20,000 < N < 100,000 (honey with low pollen content);
- Class III: 100,000 < N < 500,000 (honey with high pollen content);
- Class IV: 500,000 < N < 1,000,000 (honey with very high pollen content);
- Class V: N > 1,000,000 (honey with extremely high pollen content).

## RESULTS

### Pollen taxa from honeys of the sub-prefecture of Cechi

Melissopalynological analysis identified 59 pollen taxa across all honey samples. Fifty-four (91.53%) were identified taxa, four (6.78%) were undetermined, and one (1.69%) was a spore. The identified taxa belonged to 24 botanical families. The families with the largest number of pollen taxa were Fabaceae (9 species, 16.67%), Apocynaceae, and Combretaceae, each with 5 species (9.26%) (Fig. 1). Pollen taxa such as *Elaeis guinensis* (Arecaceae), *Cyperaceae-type* (Cyperaceae), *Bridelia micrantha* (Euphorbiaceae), and *Verbenaceae-type* (Verbenaceae) were found in all analyzed honey samples, representing a 100% occurrence rate. These are followed by taxa such as: *Scotellia chevaleri* (Flacourtiaceae), *Daniellia oliveri* (Fabaceae), *Caesalpiniaceae-type* (Fabaceae), *Lamiaceae-type* (Lamiaceae), and *Rauvolfia vomitoria* (Apocynaceae), found in four out of five (4/5) honey samples, representing an occurrence rate of 80%. Thirteen taxa are found in three out of five (3/5) honey samples, representing an occurrence rate of 60%. These are: *Poaceae-type* (Poaceae), *Digitaria-type* (Poaceae), *Syzygium-type* (Myrtaceae), *Malvaceae-type* (Malvaceae), *Bombax buonopozense* (Malvaceae), *Senna siamea* (Fabaceae), *Acacia pennata* (Fabaceae), *Pterygota macrocarpa* (Combretaceae), *Combretum gradiflorum* (Combretaceae), *Apocynaceae-type* (Apocynaceae), *Alstonia boonei* (Apocynaceae), *Mangifera indica* (Anacardiaceae) and *Lannea-type* (Anacardiaceae). The remaining taxa are taxa found in two honey samples out of five (2/5), representing an appearance rate of 40%, and those present in one honey sample out of five (1/5), representing an appearance rate of 20%.

**Figure 1:**
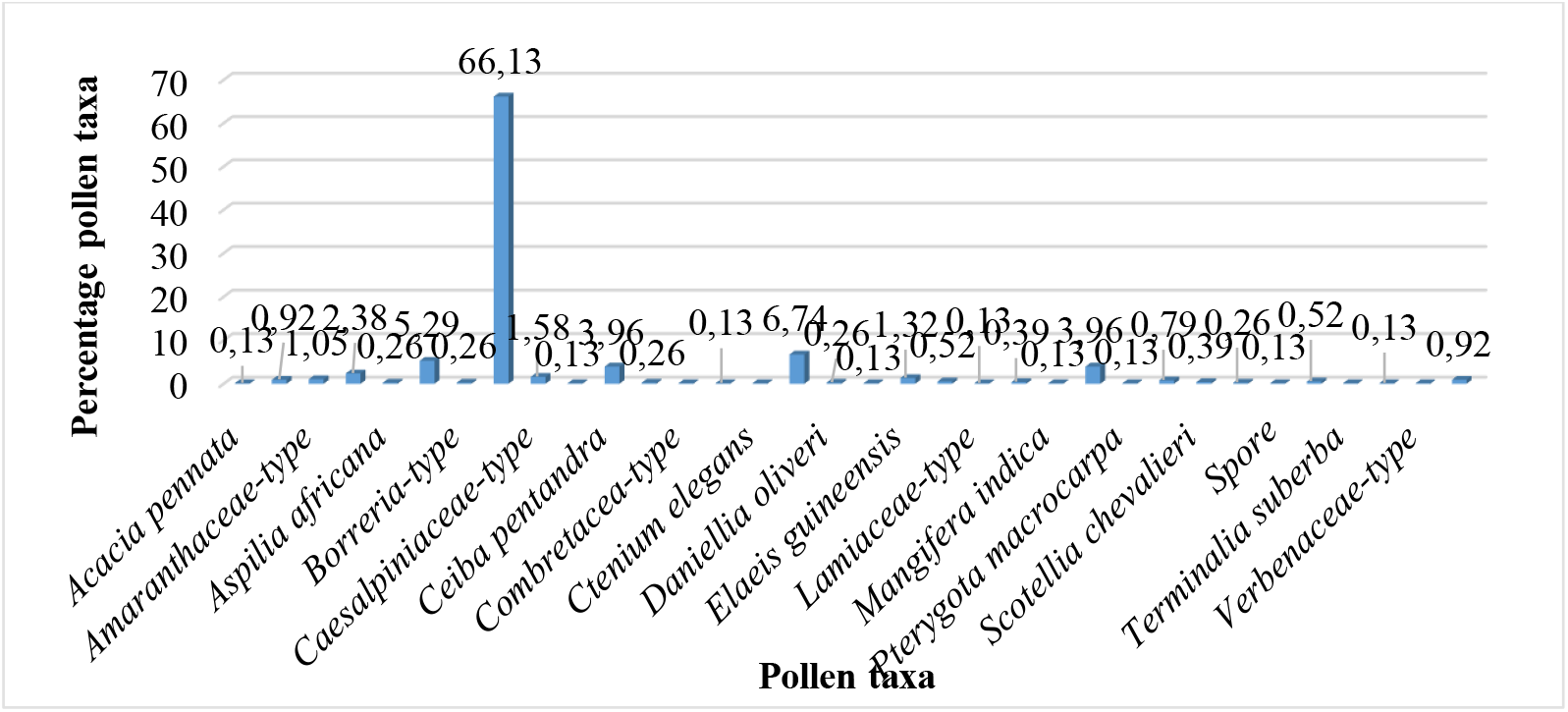
Pollen spectrum of the honey sample from the Cechi Nature Reserve (E3)

The pollen plates are presented following the bibliographical references in the appendix.

### Floral Names of Honeys from the Study Area

The pollen taxon richness of the studied honey samples ranged from 18 to 34 taxa. Honey sample (E3) from modern hives located in the Boua M’po Nature Reserve contained the highest number of pollen taxa, with 31 identified taxa, some unidentified taxa, and spores (Fig. 1). The dominant pollen taxon in this honey was Bridelia micrantha (Euphorbiaceae), representing 66.13% of the pollen grains. Four other important isolated pollen taxa were found, with frequencies ranging from 3% to 16%: *Cyperaceae-type* (Cyperaceae), *Bombax buonopozense* (Malvaceae), *Ceiba pentandra* (Malvaceae), and *Poaceae-type* (Poaceae). Other pollen taxa and spores are rare isolated pollens with frequencies of less than 3%.

The honey sample (E4) from Mitichi follows with 31 pollen taxa (Fig. 2). Two of its pollen taxa are accompanying pollen grains with frequencies ranging from 16% to 45%: *Elaeis guineensis* (Arecaceae) and *Scotellia chevalieri* (Flacourtiaceae). Three of its pollen taxa are significant isolated pollen grains with frequencies ranging from 3% to 16%: *Alstonia boonei* (Apocynaceae), *Bridelia micrantha* (Euphorbiaceae), and *Verbenaceae-type* (Verbenaceae). Its other pollen taxa are rare isolated pollen grains with frequencies below 3%.

**Figure 2.**
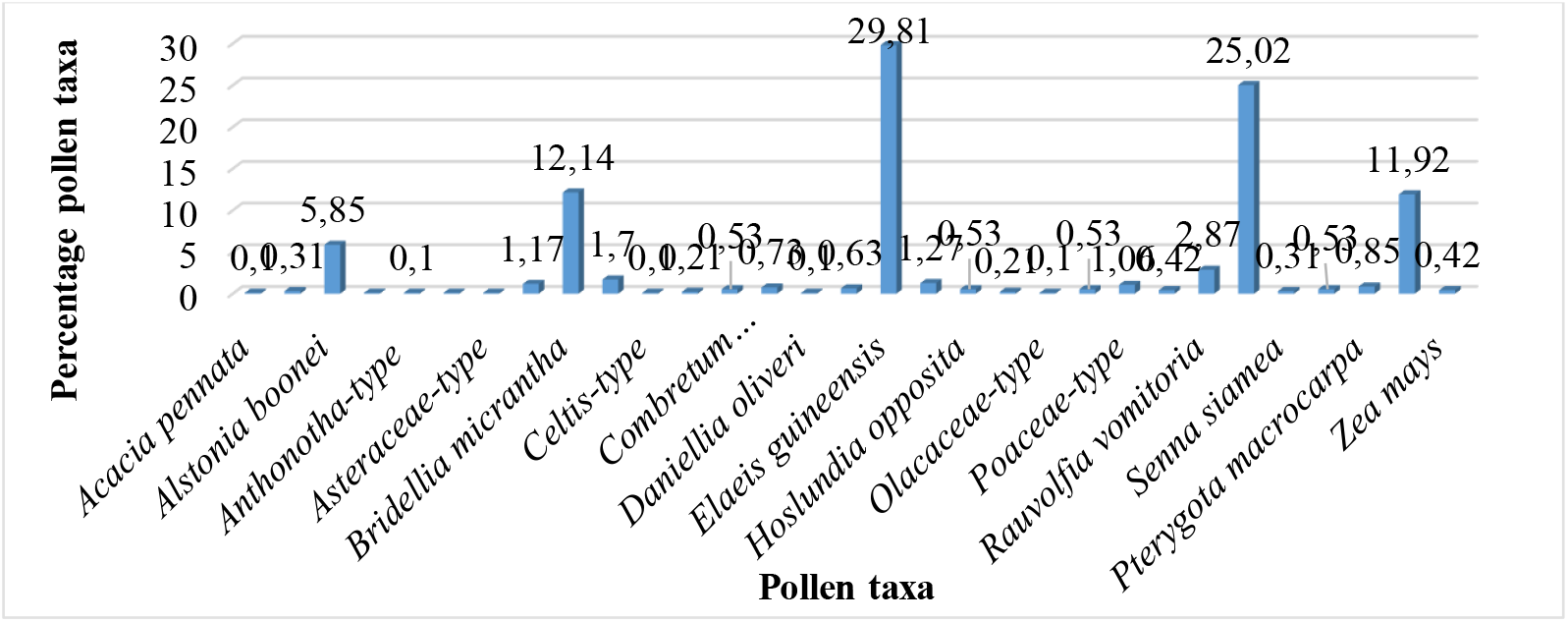
Pollen spectrum of the honey sample from Mitichi (E4)

The honey sample (E5) from N’guessanBlekro comes in third place with 26 pollen taxa (Fig. 3). One of its pollen taxa is an accompanying pollen grain with a frequency of 34.48%. This is *Blighia welwitschii* (Sapindaceae). Five of its pollen taxa are significant isolated pollen grains with frequencies between 3% and 16%. These are: *Mangifera indica* (Anacardiaceae), *Bridelia micrantha* (Euphorbiaceae), *Syzygium-type* (Myrtaceae), *Olacaceae-type* (Olacaceae), and *Verbenaceae-type* (Verbenaceae). Its other pollen taxa are rare isolated pollen grains with frequencies below 3%.

**Figure 3.**
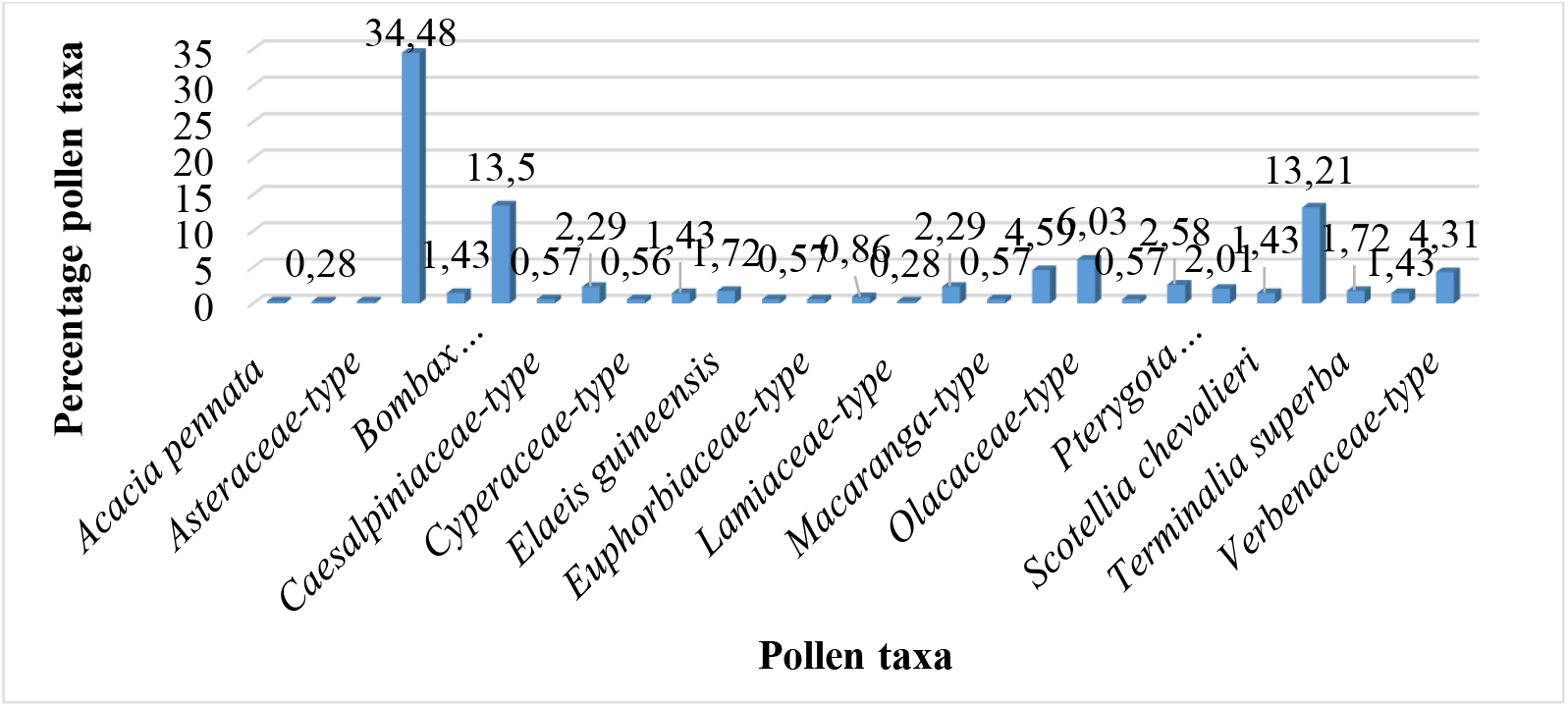
Pollen spectrum of the honey sample from N’guessanBlekro (E5)

The Allany honey sample (E1) ranks fourth with 19 identified pollen taxa and some unidentified taxa (Fig. 4).

**Figure 4:**
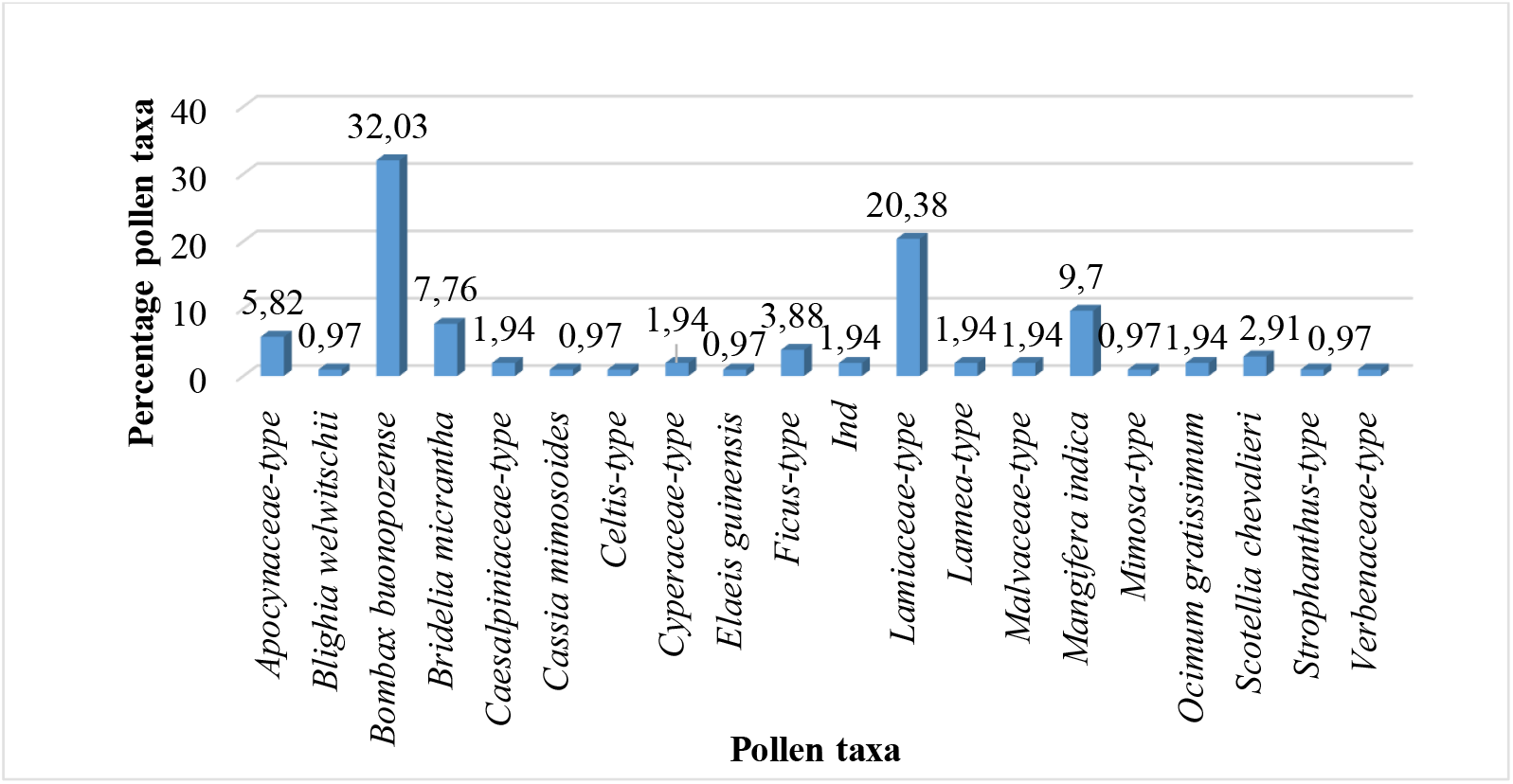
Pollen spectrum of the honey sample from Allany (E1)

Two of its pollen taxa are accompanying pollen grains with frequencies ranging from 16% to 45%: *Bombax buonopozense* (Malvaceae) and *Lamiaceae-type* (Lamiaceae). Four of its pollen taxa are significant isolated pollen grains with frequencies ranging from 3% to 16%: *Apocynaceae-type* (Apocynaceae), Bridelia micrantha (Euphorbiaceae), *Lamiaceae-type* (Lamiaceae), and *Mangifera indica* (Anacardiaceae). Its remaining pollen taxa are rare isolated pollen grains with frequencies below 3%.

The honey sample (E2) from Banguié 2 has the lowest pollen count. Sixteen pollen types were identified (Fig. 5). One of these pollen types is an accompanying pollen grain with a frequency of 41.95%. This is Lamiaceae-type. Five of these pollen types are significant isolated pollen grains with frequencies ranging from 3% to 16%. These are: Apocynaceae-type, Rauvolfia vomitoria, Bridelia micrantha, Antiaris-type, and Verbenaceae-type. The remaining pollen types are indeterminate pollen grains with frequencies below 3% (rare isolated pollen). Of all the honey samples analyzed, four did not contain dominant pollen grains (RF < 45%). Only the sample (E3) from the nature reserve contains a dominant pollen (RF = 66.13%).

**Figure 5.**
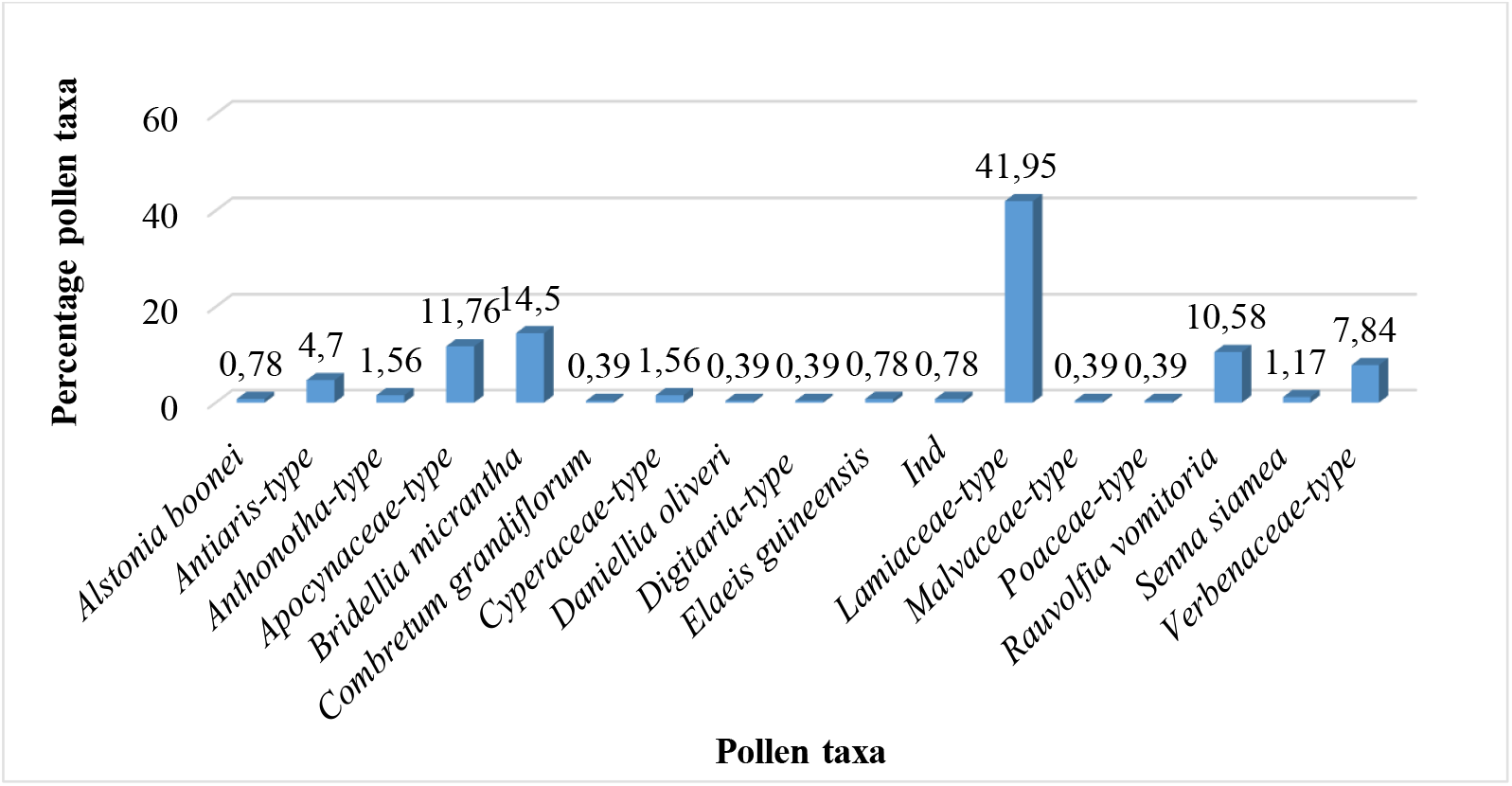
Pollen spectrum of the honey sample from Banguié 2 (E2)

### Pollen Content of Honeys from the Study Site

The pollen count in 20 µL of a microscopic slide of these honeys ranged from 309 pollen grains (E1) to 2,817 pollen grains (E4). The pollen content calculated per 10 g of the analyzed honeys ranged from 216,300 pollen grains (E1) to 1,873,305 pollen grains (E4).

The honeys studied were divided into three classes or categories according to their pollen content. The Allany honey (E1) is a Class III honey (honey rich in pollen). The Banguié 2 honey (E2): 531,675 pollen grains and the N’guessanBlekro honey (E5): 783,000 pollen grains are Class IV honeys (honey very rich in pollen). Those from the nature reserve (E3) in Boua M’po and from Mitichi (E4) are class V honeys (honey extremely rich in pollen).

## DISCUSSION

Palynological analysis of honeys is a complementary method to field inventories of melliferous plants [33, 4, 26]. It allows for the identification of plants foraged by honeybees based on the pollen taxa contained in the honey. Thus, melissopalynology makes it possible to determine the floral and geographic origins of honeys. A total of fifty-four (54) pollen taxa were identified for all 5 honey samples analyzed from the sub-prefecture of Cechi. This number of pollen taxa identified by the study is closer to the 61 pollen taxa identified by Xiao-Yan et al. [25] in 19 samples of natural honey from the central Shanxi region of northern China. Moreover, the number of pollen taxa identified by the study is greater than the number of pollen taxa identified in 13 honey samples from the wooded savannas of Togo and Benin (39 pollen taxa) according to Lobreau-Callen et al. [20].

However, it is lower than the number of pollen taxa identified by Kouassi [34] (85 pollen taxa) in 10 honey samples from Katiola; by Coulibaly [35] (206 pollen taxa) in 16 honey samples from localities in the Central (Dimbokro), Northern (Katiola, Korhogo), and Western (Danané) regions of Côte d’Ivoire; by Gnangouli bi et al. [14] (588 pollen taxa) in 20 honey samples from the Central and Northern regions of Côte d’Ivoire: Poro, Tchologo, Hambol, Bélier, and N’zi.

Furthermore, by Comlan et al. [26], (82 pollen taxa) in 4 samples of honey from localities in the Guinean zone of Togo (Danyi-Elavanyo, Igbélékoutsè-Béna, Azianfokopé-Takpla and Danyi-Akayo).

The difference in the number of pollen taxa in honeys could be explained by the number of honey samples analyzed, the diversity of nectar-producing flora flowering in the bees’ foraging area during the honey flow, or the diversity of plants foraged by the bees to produce the honey. Regarding the botanical families (Fabaceae, Apocynaceae, and Combretaceae) best represented in pollen taxa, this could be explained by the quality of the pollen of their species or by the flowering of most species during the honey flow period. Melissopalynological studies conducted worldwide, elsewhere in Africa, and in Côte d’Ivoire have also shown a dominance of taxa belonging to the Fabaceae [36, 7, 25, 26, 34] and Combretaceae [13, 14] families. The abundance of taxa from the Fabaceae and Combretaceae families could be an asset for beekeeping and honey production in the region.

The large number of pollen taxa from the Apocynaceae family in the honeys and taxa such as: *Elaeis guineensis* (Arecaceae), *Cyperaceae-type* (Cyperaceae), *Bridelia micrantha* (Euphorbiaceae) and *Verbenaceae-type* (Verbenaceae) present in all the honey samples analyzed could constitute a peculiarity or botanical signature for the honeys of the region, although the richness of pollen taxon in the honey depends on the melliferous plants surrounding the apiaries and following the foraging area of the bees.

The high pollen taxon count of the honeys, ranging from 18 to 34 taxa, with the honey from the nature reserve having the highest count, demonstrates the diversity of nectar-producing plants in the reserve and its importance for beekeeping in the area. Furthermore, beekeeping practices near protected areas (classified forests, nature reserves, and national parks) improve honey production, the sustainability of plant species, and allow local communities (beekeepers) to participate in the preservation of surrounding forests, as their beekeeping production depends on it. The pollen taxon count of the analyzed honeys is comparable to honeys from the Guinean zone of Togo (25 to 38 taxa) according to Comlan et al. [26], and to Toumodi honey (27 taxa) according to Assi et al. [13]. Moreover, the richness in pollen taxa of the studied honeys is greater than that of the honeys from the central Shanxi region in Northern China (7 to 22 taxa) according to Xiao-Yan et al. [25] and of the honeys from the Worodougou region (15 to 31 taxa) in Northern Ivory Coast according to Diomandé et al. [36].

Regarding the floral classification of the honeys, most of the analyzed honeys (4/5) do not contain dominant taxa (RF > 45%). They consist of accompanying pollen taxa (16% ≤ RF < 45%), significant isolated taxa (3% ≤ RF < 16%), and rare isolated taxa (RF < 3%). These honeys are therefore polyfloral, multifloral, or wildflower honeys. The honey sample from the reserve, with more than 45% Bridelia micrantha pollen grains (RF = 66.13%), is a monofloral or unifloral honey and could be botanically classified as *Bridelia micrantha* honey. The study area in particular, and the Ivorian forest zone in general, thus produces both polyfloral and monofloral honeys.

Regarding the pollen content of the honeys, the analyzed samples were Class III (pollen-rich honey), Class IV (very pollen-rich honey), and Class V (extremely pollen-rich honey). These are natural honeys sought after on the market due to their high pollen grain content, which attests to their quality. According to Indian legislation on the pollen grain content of commercially available honeys, high-quality honeys of natural origin must have a pollen grain content exceeding 50,000 grains [37]. The pollen content of the Class III (E1) and Class IV (E2, E5) honeys from the study area is similar to the honey from Florona (Class III) and Wongué (Class IV) in the Séguéla department [36] and to the honey from the Toumodi department [13] (Class III). These honeys are richer in pollen grains than the honeys from Bobi (class II) in the department of Séguéla and from the commercial center of the city of Daloa (class I) according to Diomandé et al. [36].

The Class V honeys identified by the study: E4 and E3 from the reserve (honeys extremely rich in pollen) are richer in pollen than the honeys studied from other regions of Côte d’Ivoire. This is supported by the work of Coulibaly [35], Diomandé et al. [36], and Assi et al. [13], as well as honeys from Northern China according to Xiao-Yan et al. [25]. Furthermore, the pollen content of honeys is thought to be linked to the quantity of pollen-producing plants available in the environment, plants that flowered during the honey flow, or to the different honey extraction techniques. According to Randrianarivelo [7], honeys very rich in pollen (Class IV) and honeys extremely rich in pollen (Class V) are pressed honeys.

## CONCLUSION

A melissopalynological study conducted on honeys from the sub-prefecture of Cechi, in the southern forest region of Côte d’Ivoire, identified 54 out of 59 recorded pollen taxa, representing a rate of 91.53%. The Fabaceae, Apocynaceae, and Combretaceae families are the most represented among melliferous plant species in the region. Twenty-two pollen taxa constitute the main pollen taxa in the area’s honeys, with an occurrence rate of over 50%. The honeys from the area are of two types, based on their taxonomic content: monofloral and polyfloral. The majority of honeys are produced by bees from the flowers of several plant species (polyfloral honeys). Monofloral honey, with over 45% of its pollen grains being *Bridelia micrantha* (66.13%), is the honey of the reserve. This type of honey could bear the botanical name of the species.

All the honeys analyzed have a high pollen content, which justifies their good quality and natural origin from a palynological point of view. These are honeys rich, very rich, and extremely rich in pollen. The significant number of Apocynaceae pollen taxa in the honeys, and taxa such as *Elaeis guineensis* (Arecaceae), *Cyperaceae-type* (Cyperaceae), *Bridelia micrantha* (Euphorbiaceae), and *Verbenaceae-type* (Verbenaceae) present in all the honey samples analyzed, could constitute a particularity or botanical signature for the honeys of the region.

## ACKNOWLEDGEMENTS

We would like to thank all the directors of the laboratory at the Faculty of Pharmaceutical and Biological Sciences of Félix Houphouët-Boigny University (UFHB), who allowed us to conduct our various experiments in their laboratory and provided us with all the necessary equipment.

We also thank the Botany Research Teaching Unit and the Biosciences Faculty of UFHB, to which we are affiliated.

---

**Appendix 1:**
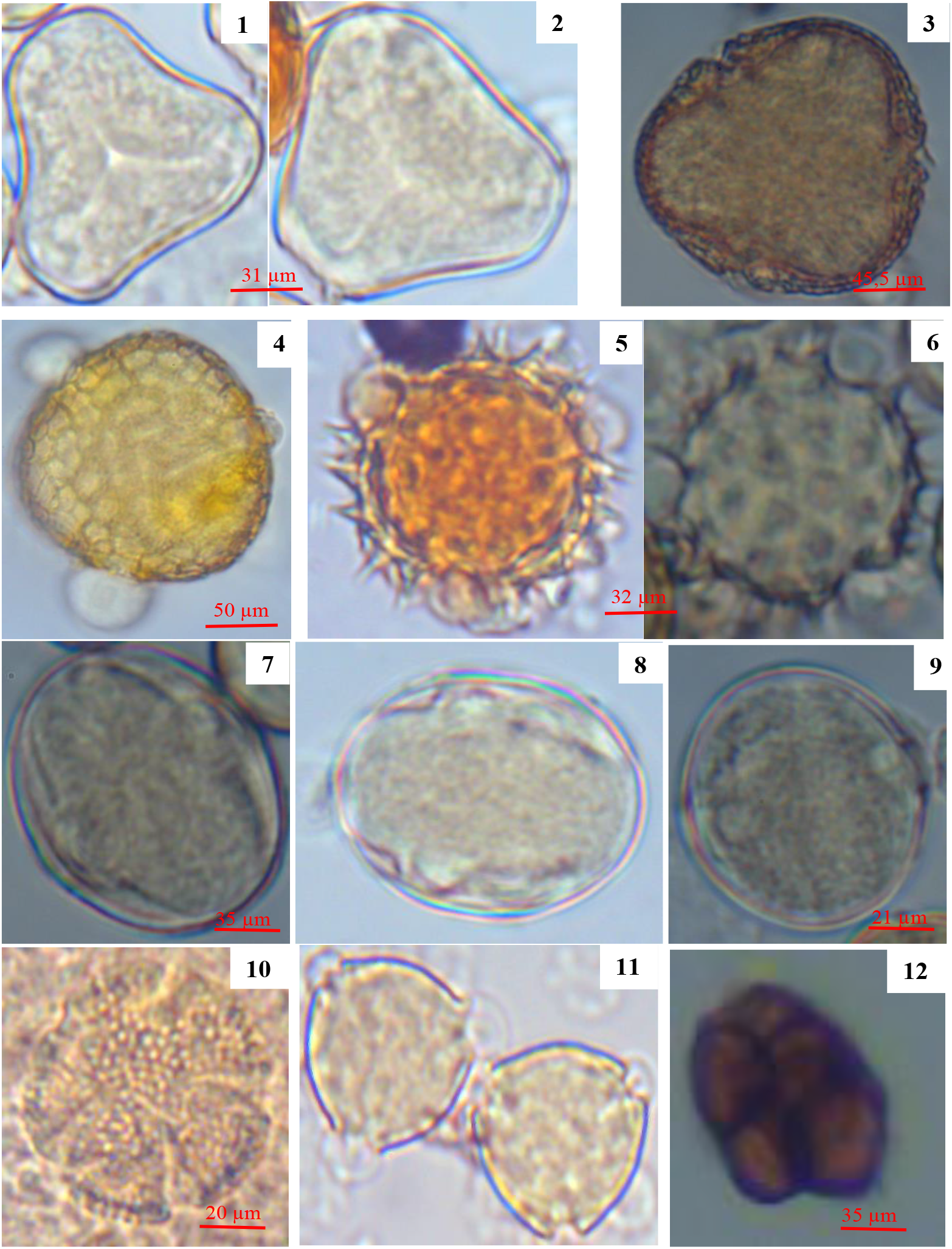
Pollen grains from honey producing taxa in the sub-prefecture of Cechi. 1,2: *Elaeis guineensis* (Arecaceae); 3: *Bombax buonopozense* (Malvaceae); 4: *Ceiba pentandra* (Malvaceae); 5,6: *Aspilia africana* (Asteraceae); 7: *Combretum grandiflorum* (Combretaceae); 8: *Pterygota macrocarpa* (Malvaceae); 9: *Terminalia superba* (Combretaceae); 10: *Ocimum gratissimum* (Lamiaceae); 11: *Alstonia boonei* (Apocynaceae); 12: *Acacia pennata* (Fabaceae)

**Appendix 2:**
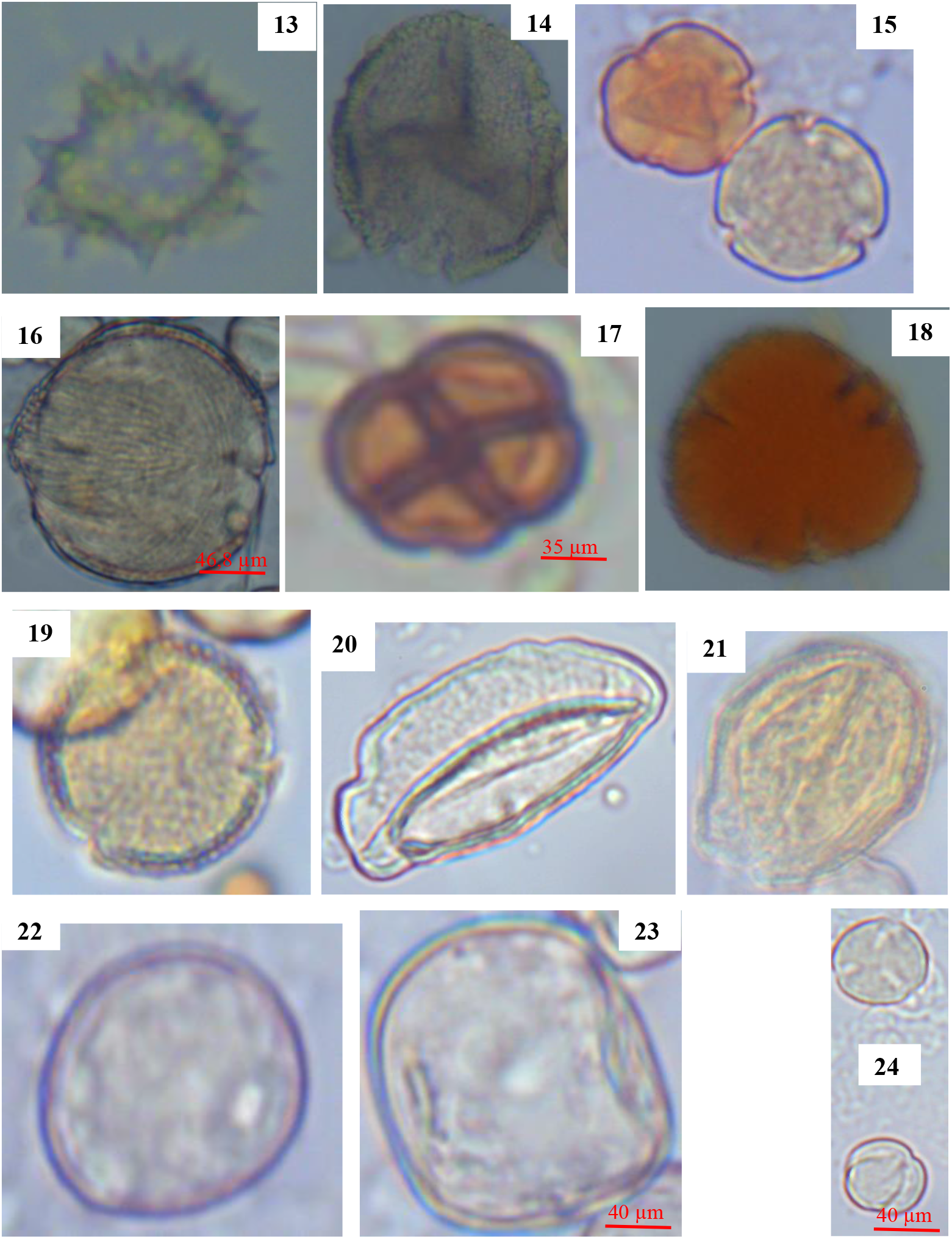
Pollen grains from honey producing taxa in the sub-prefecture of Cechi. 13: *Asteraceae-type* (Asteraceae); 14: *Euphorbiaceae-type* (Euphorbiaceae); 15: *Alafia-type* (Apocynaceae); 16: *Rauvolfia vomitoria* (Apocynaceae); 17: *Albizia-type* (Fabaceae); 18: *Malvaceae-type* (Malvaceae); 19: *Pterygota aubrevillei* (Sterculiaceae); 20: *Cyperaceae-type* (Cyperaceae); 21: *Cassia mimosoides* (Fabaceae); 22: *Poaceae-type* (Poaceae); 23: *Digitaria-type* (Poaceae); 24: *Bridelia micrantha* (Euphorbiaceae)

**Appendix 3:**
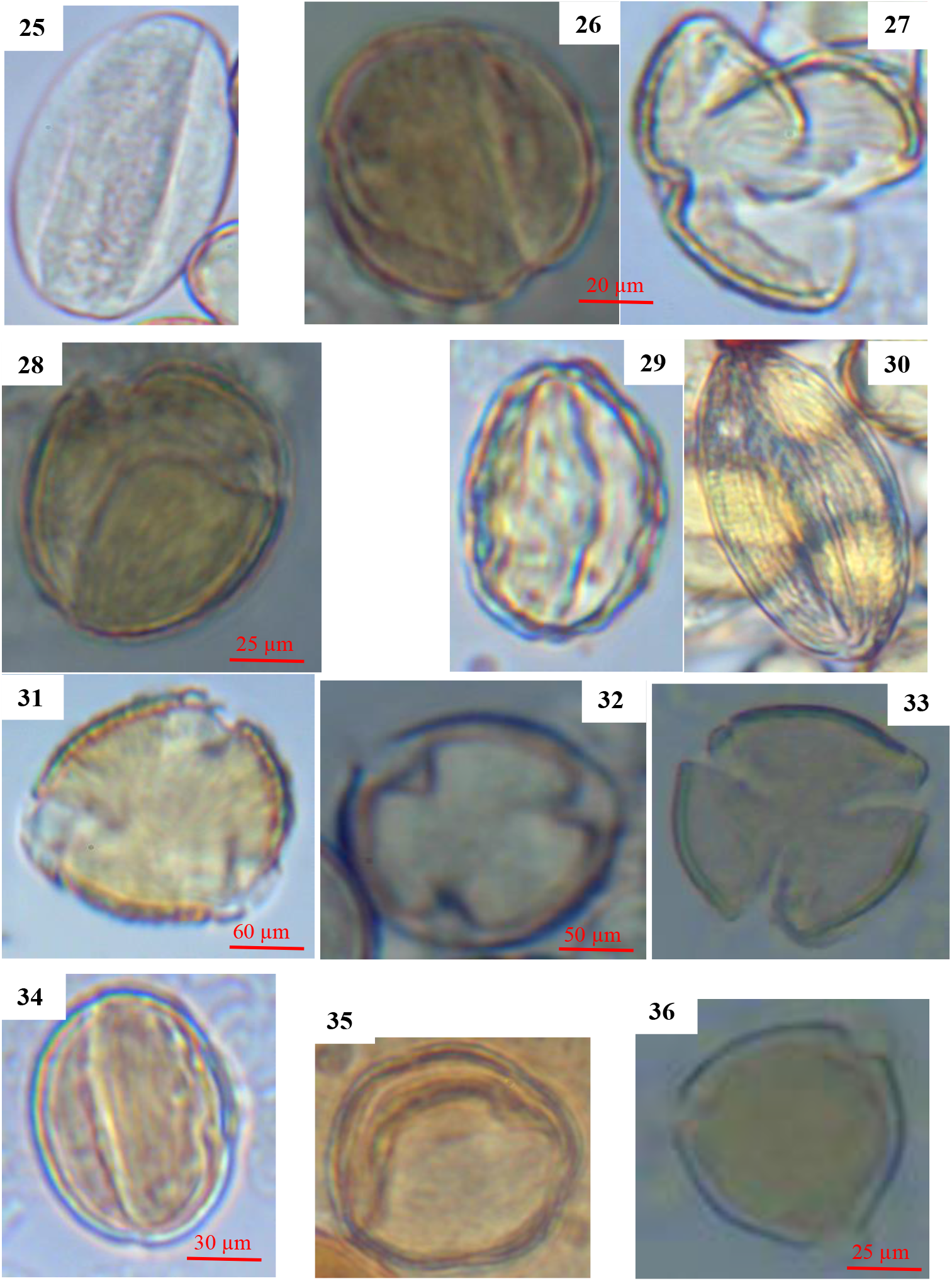
Pollen grains from honey producing taxa in the sub-prefecture of Cechi. 25: *Lamiaceae-type* (Lamiaceae); 26, 27: *Blighia welwitschii* (Sapindaceae); 28: *Lannea-type* (Anacardiaceae); 29, 30: *Anthonotha-type* (Annonaceae); 31: *Isoberlinia doka* (Fabaceae); 32: *Daniellia oliveri* (Fabaceae); 33: *Clerodendrum-type* (Verbenaceae); 34: *Verbenaceae-type* (Verbenaceae); 35: *Celtis-type* (Ulmaceae); 36: *Mangifera indica* (Anacardiaceae)

**Appendix 4:**
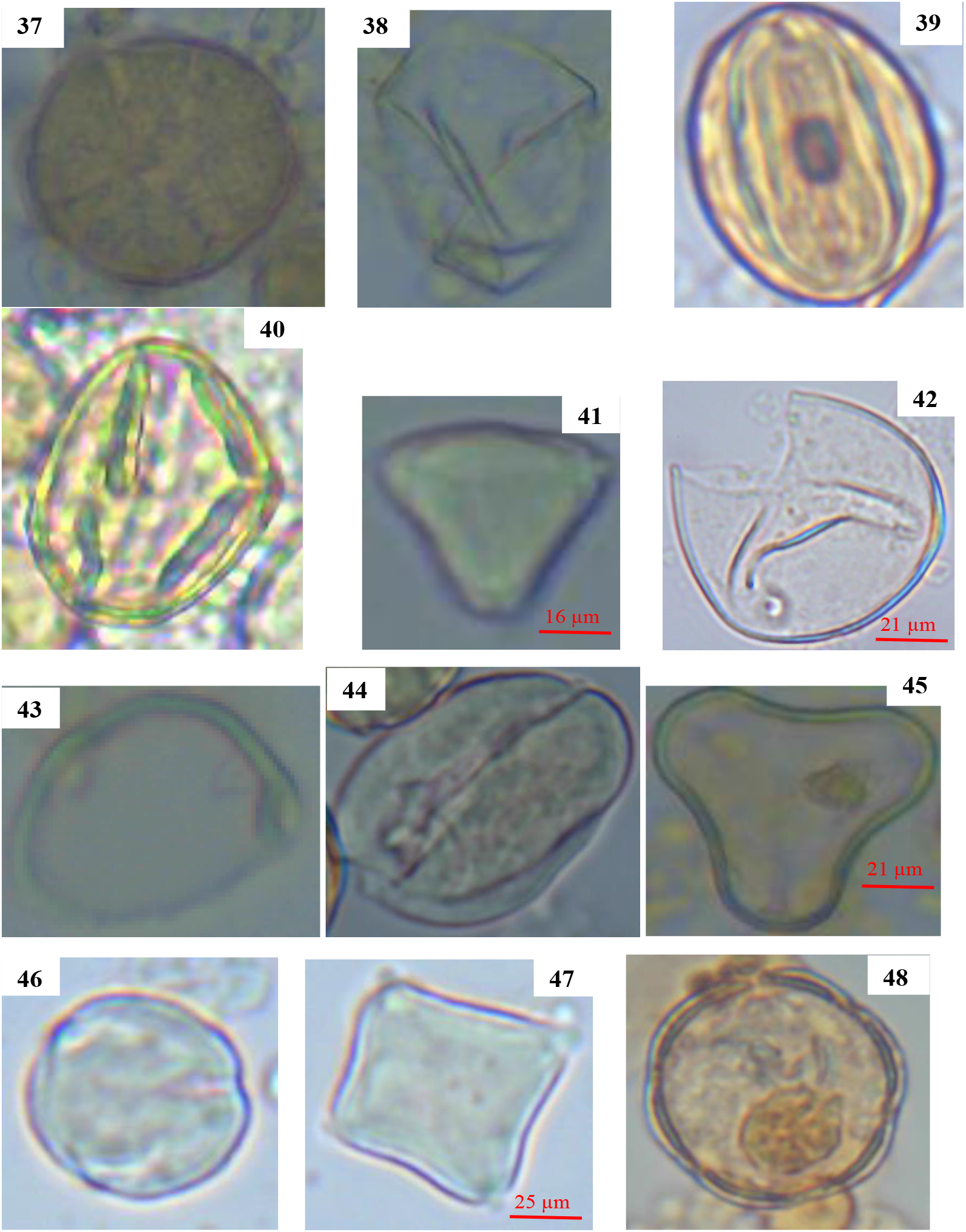
Pollen grains from honey producing taxa in the sub-prefecture of Cechi. 37: *Entandrophragma-type* (Meliaceae); 38: *Zea mays* (Poaceae); 39: *Terminalia-type* (Sterculiaceae); 40: *Caesalpiniaceae-type* (Fabaceae); 41: *Syzygium-type* (Myrtaceae); 42: *Strophanthus-type* (Apocynaceae); 43: *Ficus-type* (Moraceae); 44: *Annona-type* (Annonaceae); 45: *Macaranga-type* (Euphorbiaceae);46: *Senna siamea* (Fabaceae); 47: *Olacaceae-type* (Olacaceae); 48: *Lamiaceae-type* (Lamiaceae)

**Appendix 5:**
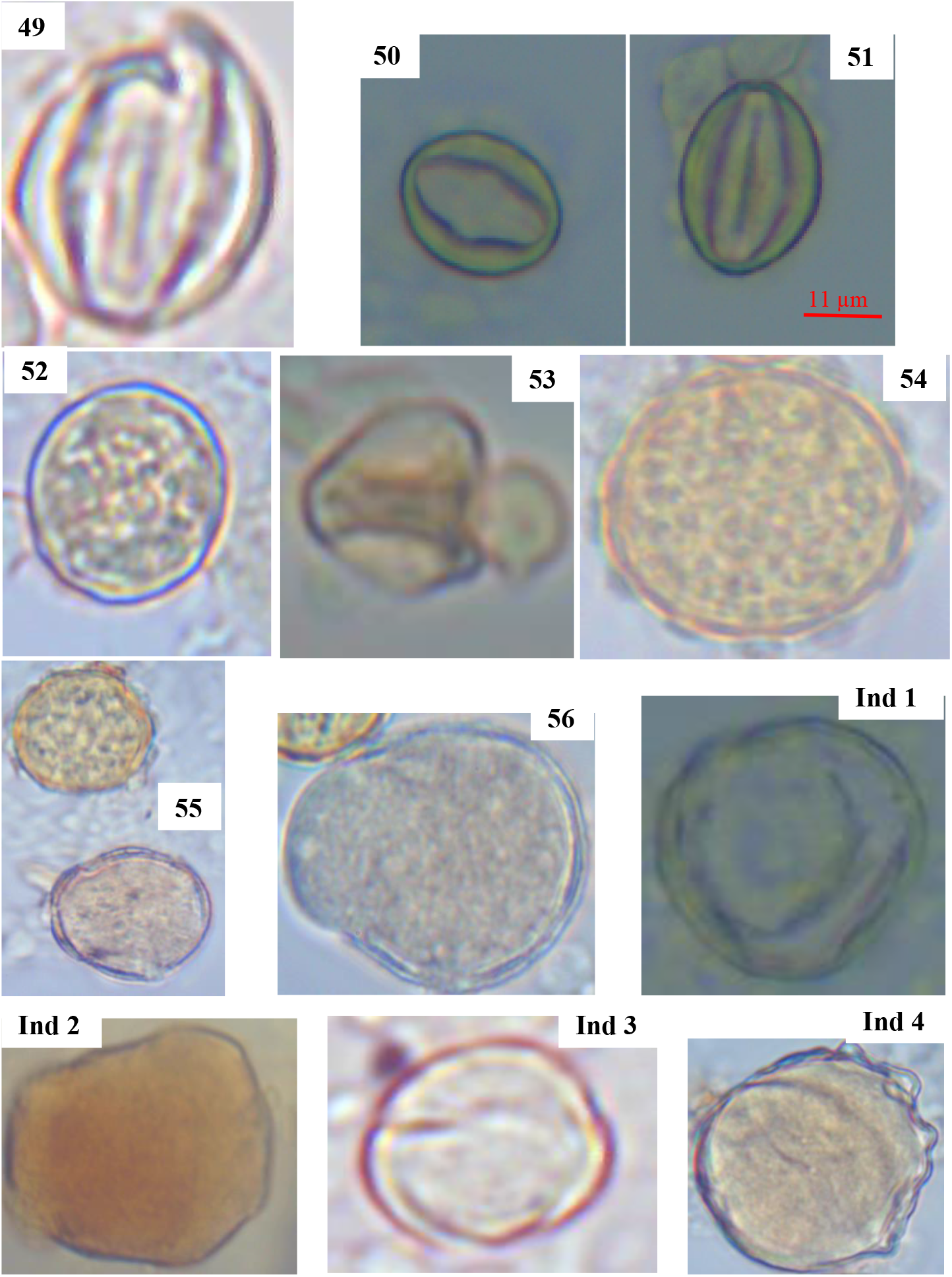
Pollen grains from honey producing taxa in the sub-prefecture of Cechi. 49: *Combretaceae-type* (Combretaceae); 50, 51: *Scotellia chevalieri* (Flacourtiaceae); 52: *Amaranthaceae-type* (Amaranthaceae); 53: *Mimosa-type* (Fabaceae); 54: *Borreria-type* (Rubiaceae); 55: *Convolvulaceae-type* (Convolvulaceae); 56: *Apocynaceae-type* (Apocynaceae), Ind: Indéterminé

